# Free Energy Landscape and Conformational Kinetics of Hoogsteen Base-Pairing in DNA vs RNA

**DOI:** 10.1101/2020.01.09.868018

**Authors:** D. Ray, I. Andricioaei

## Abstract

Genetic information is encoded in the DNA double helix which, in its physiological milieu, is characterized by the iconical Watson-Crick nucleobase pairing. Recent NMR relaxation experiments revealed the transient presence of an alternative, Hoogsteen base pairing pattern in naked DNA duplexes and estimated its relative stability and lifetime. In contrast, HG transitions in RNA were not observed. Understanding Hoogsteen (HG) base pairing is important because the underlying "breathing" can modulate significantly DNA/RNA recognition by proteins. However, a detailed mechanistic insight into the transition pathways and kinetics is still missing. We performed enhanced sampling simulation (with combined metadynamics and adaptive force bias method) and Markov State modeling to obtain accurate free energy, kinetics and the intermediates in the transition pathway between WC and HG base pair for both naked B-DNA and A-RNA duplexes. The Markov state model constructed from our unbiased MD simulation data revealed previously unknown complex extra-helical intermediates in this seemingly simple process of base pair conformation switching in B-DNA. Extending our calculation to A-RNA, for which HG base pair is not observed experimentally, resulted in relatively unstable single hydrogen bonded distorted Hoogsteen like base pair. Unlike B-DNA the transition pathway primarily involved base paired and intra-helical intermediates with transition timescales much higher than that of B-DNA. The seemingly obvious flip-over reaction coordinate, i.e., the glycosidic torsion angle is unable to resolve the intermediates; so a multidimensional picture, involving backbone dihedral angles and distance between atoms participating in hydrogen bonds, is required to gain insight into the molecular mechanism.

**SIGNIFICANCE:** Formation of unconventional Hoogsteen (HG) base pairing is an important problem in DNA biophysics owing to its key role in facilitating the binding of DNA repairing enzymes, proteins and drugs to damaged DNA. X-ray crystallography and NMR relaxation experiments revealed the presence of HG base pair in naked DNA duplex and protein-DNA complex but no HG base pair was observed in RNA. Molecular dynamics simulations could reproduce the experimental free energy cost of HG base pairing in DNA although a detailed mechanistic insight is still missing. We performed enhanced sampling simulation and Markov state modeling to obtain accurate free energy, kinetics and the intermediates in the transition pathway between WC and HG base pair for both B-DNA and A-RNA.

## INTRODUCTION

The one-dimensional genetic information encoded in the sequence of DNA base pairs is intrinsically related to its three-dimensional structure described by the iconical Watson-Crick (WC) helix (1), with its specific hydrogen-bonding pattern between purine and pyrimidine (A-T, C-G) complementary nucleobases.

Recently, a series of exciting NMR relaxation studies (2) on free DNA in solution have shown that the predominant WC base-pairing pattern between A-T and G-C nucleotides is in dynamic equilibrium with a less common and transient (*µ*s-ms lifetime) Hoogsteen (HG) form (3), where the purine nucleotide in the base pair flips over. The WC to HG base pair transition, a form of “DNA breathing” (4), hints to the existence of a “secondary genetic code” signaled by base pair flipping, apart from the primary encoding in the nucleotide sequence. Crystallographic experiments have reported the presence of Hoogsteen base pairs only in specific contexts, such as when DNA binds to proteins like p53 tumor suppressor protein (5) or integration host factor (IHF) in *E. coli* (6). Detailed biochemical studies showed that HG base pairing plays a key role in circumventing interference in DNA replication mechanism by DNA polymerase by avoiding appearance of lesions (7, 8). In p53 binding sequence HG conformation exposes negative charged regions of nucleic acid to the positive arginine residue of the protein (5) leading to stability of the complex and playing crucial role in DNA recognition (9, 10). Examples of specific binding to DNA guided by HG base pair extends beyond proteins and even includes small molecules and drugs, which in turn facilitates its recognition by proteins and enzymes (11, 12). In a nutshell, HG base pairing therefore expands the structural and functional repertoire of duplex DNA beyond what can be achieved by Watson-Crick base-pairing alone. For example, HG base pairs are recognized by DNA repairing enzymes resulting in selective binding in the damaged regions of DNA, rich in syn-anti configuration over the anti-anti WC base pair rich normal DNA. Thus, apart from providing structural integrity to the damaged and deformed DNA, HG base pairs assist in the DNA repair mechanism (13).

Apart from multiple appearance of HG base pairing in damaged or distorted DNA (14–16) several studies have confirmed presence of Hoogsteen conformation in undistorted DNA duplex (17, 18); the first X-ray crystallographic evidence of Hoogsteen base pairing in a simple stretch of DNA sequence composed exclusively of A-T base pairs was provided by Abrescia and coworkers (19). Early computational work by Gould and Kollman had showed that HG base pairing between adenine and thymine nucleobases (i.e., without sugar-phosphate backbone) in vacuum was energetically more favourable (~ 1 kcal/mol) than its WC counterpart (20) in agreement with the structure in crystalline environment (3). Subsequent ab-initio quantum chemistry calculations for gas phase A-T base pairs have painted a more complex picture, showing that neither HG or WC forms are the global minimum (21).

Furthermore, NMR studies have found that N-methylation of adenine favors *syn* HG base pairing over the *anti* WC form by preventing formation of a WC specific hydrogen bond by inducing steric clash (16). NMR chemical shift perturbation, NOESY experiments and constant pH MD simulation show that the HG base pairing among G-C base pairs is ~20 times less abundant than that in A-T base pairs arguably because a protonation of cytosine is necessary for the transition to HG form (11, 22). Experiments of bare DNA duplex like A-T rich A6-DNA segment have shown presence of Hoogsteen base pair in the A16-T9 position from carbon *R*_1ρ_ relaxation dispersion NMR signal (2). Such experiments on DNA bound to base intercalating drugs like echinomycin and actinomycin showed formation of Hoogsteen base pair (23).

Since RNA duplexes also involve Watson-Crick like base pairs, it was natural to look for the existence of Hoogsteen transitions there, as well. However, HG base pairing was not observed in NMR relaxation experiment in a common A-form RNA hairpin with same sequence as A6-DNA. In contrast to DNA N-methylation of RNA adenine base to induce HG base pair formation resulted in base pair melting and consequent dissociation of the RNA helical region (24). In the present paper, we perform a computational comparison of the free energy psurface and transition paths of DNA vs RNA WC to HG transitions, seeking to understand the relative stability of the HG state and the factors that modulate the dynamical equilibrium in DNA vs RNA.

The first report (2) of the NMR measured WC-HG dynamic equilibrium also included our preliminary work on searching pathways of conformational transition using a biased molecular dynamics simulation. Further computational work from our group and others (12, 23–26) have since uncovered several other pieces of information, including energetic details, transition pathways and kinetics of the transition between WC and HG base pairs in naked A-T rich DNA helical duplexes. The paths predicted in our initial study were also recently sampled by Vreede et al.(27), who have calculated the rate constants of back and forth transitions between WC and HG base pairs using Transition Path Sampling (TPS) (28).

Experimental estimates put the WC-HG transition in DNA on a timescale of 50-250 ms (2). Therefore, direct simulation of Hoogsteen base pair formation in explicit water for a biologically relevant system is still beyond the reach of present day computational power. In previous work, we showed that formation of Hoogsteen base pair can be achieved in molecular dynamics simulation by applying an artificial biasing force on the the glycosidic torsion angle for free DNA (2, 24) and echinomycin-bound DNA (23).

Enhanced sampling methods like umbrella sampling (29) and metadynamics (30, 31) can be utilized to obtain the free energy surface along slow coordinates. For example, Yang et. al. have calculated the free energy landscape of DNA breathing motion for a A-T rich duplex DNA segment from umbrella sampling (25) and multiple walker well tempered metadynamics simulation(26). Even for a short DNA segment, it required 6-40 *µ*s long enhanced sampling simulations to obtain a converged physically meaningful free energy surface.

Adaptive biasing force (ABF) (32) has recently become a good alternative to the traditional free energy sampling methods (33) for a wide range of biophysical problems including membrane permeation (34), ion transport (35), ligand diffusion in protein (36) and binding free energy calculation (37).

Its recent variant named meta-eABF uses metadynamics in conjunction with extended system ABF (eABF) (38) to accelerate the sampling of the transition coordinate (39). By simultaneously depositing Gaussian bias and applying force against the potential gradient this method has been shown to be capable of sampling configurational space and predicting reasonably accurate free energy surface in orders of magnitude less simulation time (39).

We have used meta-eABF method to calculate the potential mean force (PMF) for the WC to HG transition for the duplex of A6-DNA fragment and also for A6-RNA hairpin, a synthetic construct with the same base sequence used by Zhou et al. in their experimental studies (24). Unlike A6-DNA (25, 26) there is no report of the free energy profile of the WC to HG base pair transition in A6-RNA in the literature. Moreover, revisiting DNA HG base pairing with the fast converging PMF calculation technique opens up the possibility to apply this technique in future for a large scale comparison of thermodynamics of HG base pairing in various protein and drug bound DNA complexes. Still, in the conventional PMF calculation the choice of reaction coordinate is relatively arbitrary as other collective variables might as well contribute to the conformational transition. Apart from the torsion angles of DNA (2, 25, 26), H-bond donor acceptor distances can also be considered as a viable reaction coordinate. The conformation of the thymine base is also not included in the current torsion angle based description.

Markov state models (MSM) have shown promises of alleviating this problem in molecular biophysics and found wide range of application ranging from protein conformational transitions (40, 41) to ligand unbinding (42). Construction of an MSM does not require predefined reaction coordinate and they can include all the structural information of the molecule. Also, time lagged independent component analysis (TICA) (43, 44) can be used to construct combinations of coordinate features to capture the slowest motions of the system. Provided sufficient back and forth transition sampling between clusters, it can calculate mean first passage times with reasonable accuracy (40). Therefore, we aimmed to obtain the kinetics of HG base pairing from MSM as an added analysis. Moreover, MSM can elucidate various meta-stable states in between the WC and HG conformations, which modulate base pair flipping transitions. We have constructed MSMs for both the DNA and the RNA system in order to identify the best collective variables to describe the process, construct a reaction-coordinate free free energy profile, calculate the kinetics of the process, and dissect the pathways of transition by following the probability flux through various meta-stable states.

## METHODS

### System preparation and equilibration

All input files were generated using the CHARMM-GUI web server (45, 46) and VMD (47). All simulations were performed using the NAMD 2.12 package (48) on GPUs. The NMR structure of A6-DNA (PDB ID: 5UZF) (49) was used as the starting structure for the DNA. This is a 12 base pair long DNA fragment with 6 consecutive A-T base pairs. Experimental studies have observed presence of Hoogsteen base pairing in the A16-T9 base pair, which is one of the two G-C facing A-T base pairs in this DNA fragment (2). For A6-RNA the initial structure was created using the make-NA server (50). The A6-RNA is a A-form of RNA with the same sequence as A6-DNA and at one end the two strands are stitched together with an UUCG loop (24). Both the structures were solvated using TIP3P water (51) in a rectangular solvation box with 17Å of water padding in each direction. A sufficient number of sodium ions were added to the system to neutralize the negative charge in the nucleic acid. The CHARMM36 force field for nucleic acids (52) was used in the current study. The starting structures for both DNA and RNA had all the base pairs in common Watson Crick conformation. The solvated nucleic acids were energy minimized for 50000 steps using conjugate gradient algorithm. The systems were gradually heated to 298 K temperature at a rate of 1K/ps in NVE ensemble with harmonic constraints of 3 kcal/molÅ^2^ on the nucleic acid heavy atoms. Then the constraints were removed in steps of 0.5 kcal/molÅ^2^ per 200 ps. Each system was further equilibrated for 3 ns in an NVE ensemble and 10 ns in an NPT ensemble with a small harmonic restraint on the terminal base pairs, with force constants of 0.1 kcal/molÅ^2^ and 0.05 kcal/molÅ^2^ respectively. The terminal base pairs were continually restrained with a 0.05 kcal/molÅ^2^ force constant for all production runs. The temperature was kept constant at 298 K using a Langevin thermostat with coupling constant 1 ps^−1^ and the pressure was kept constant at 1 atm using the Nosé-Hoover Langevin piston with an oscillation period of 100 fs and damping timescale of 50 fs (53, 54).

### Enhanced sampling simulation

The newly developed meta-eABF method (39) implemented in NAMD 2.12 through the colvars module (55) has been used to obtain the potential of mean force. The glycosidic angle *χ* and the pseudo-dihedral angle Θ of the A16-T9 base pair were chosen as order parameters following earlier studies (2, 23, 26, 56). The *χ* angle denotes torsion of the adenine base with respect to the deoxyribose sugar (defined as the dihedral angle between *O*4′ − *C*1′ − *N*9 − *C*4), while the Θ angle captures the flip-out motion of adenine, which leads to extra-helical conformations through base pair breaking (26). The collective variable space was spanned from −180° to 180° with a 5° bin width for both the dihedral angles. For both systems, the meta-eABF simulation was started from the final structure from the NPT equilibration and continued until all the bins were sufficiently explored and the rms difference of the PMF were converged. Gaussian biases of height 0.06 kcal/mol and width = 3 colvar units were deposited at every 2 ps for metadynamics. ABF bias was applied against the average local force only after 1000 samples were obtained at a given bin. We obtained a converged PMF within 0.2 *µ*s of simulation. The one-dimensional PMF along the glycosidic angle **χ** was obtained using Boltzmann measure integration (25)

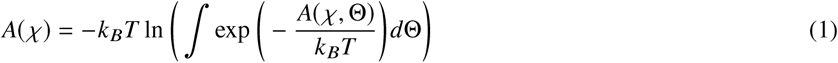

with *k*_*B*_ the Boltzmann constant, *T* the temperature, and *A χ*, Θ the 2D PMF along *χ* and Θ coordinates. The PMF along Θ was computed similarly for both DNA and RNA.

### Unbiased Simulation and Markov State Modeling

To elucidate the kinetics and the pathways of formation of Hoogsteen base pairing, Markov State Models (MSM) were built for both DNA and RNA using the PYEMMA2 package (57). A total of 45 unbiased trajectories were initiated from different starting points chosen from the biased meta-eABF trajectory. Each of them were propagated for 60 ns, resulting in 2.7 µ*s* of total trajectory data. The first 2 ns of each of the trajectories were considered as equilibration and discarded.

Total ~ 5.2 µ*s* of unbiased molecular dynamics data were used to generate the MSMs for DNA and RNA system. Trajectory snapshots were saved every 10 ps. Dimensionality reduction of the data has been performed by time-lagged independent component analysis (TICA) (43, 44) on the backbone and sugar dihedral angles (mentioned in the work of Zhou et al. (58)), the glycosidic dihedral angle the distances between hydrogen bond forming heavy atoms. Variational approach to Markov processes-2 (VAMP-2) scores (59) were calculated to estimate the appropriate lag time for TICA analysis. The trajectories were projected onto the minimum number of TICA coordinates which cover 95% of the kinetic variance. A lag time of 2 ns was chosen for both systems, which produced 5 TICA coordinates for DNA and 6 TICA coordinates for RNA. The entire simulation data was clustered into 400 clusters for DNA and 200 clusters for RNA using a k-means clustering algorithm. MSM was constructed with a lag time of 2 ns for DNA and 7 ns for RNA system. Implied timescale and Champman-Kolmogorov analyses (60) were performed to check the validity of the constructed MSM (see Figure S5–S9 in Supporting Information). The clusters were coarse grained into a few meta-stable states using the Perron-cluster cluster analysis algorithm (PCCA) (61). Kinetics and transition probabilities between the clusters were calculated using transition path theory (62, 63). The results were projected onto the coarse grained space of meta-stable states to calculate the mean first passage time (MFPT) and flux in between each pair of meta-stable states. Four sets of clusterization were performed for DNA and RNA system, and the mean and standard deviation of the first passage times were calculated.

## RESULTS AND DISCUSSION

### Free energy surfaces from meta-eABF

Unlike A6-DNA, the RNA hairpin with the same sequence did not show HG base pairing in NMR relaxation experiments (24). Forcing HG base pairing by methylation of adenine base resulted in melting of the RNA helix both in experiment and in MD simulation (24). For regular duplex RNA (un-methylated), the absence of the experimental NMR signal characteristic of HG base pairing could mean either that the HG base pair is thermodynamically unstable or that it is meta-stable, but the population and timescale fall outside the detection window of the NMR apparatus. A definitive answer to the question of HG stability in RNA can be addressed by computing the free energy landscape of base pairing conformational changes.

At first sight, the relevant degrees of freedom for the free energy landscape should contain a measure of flip-over and and flip out angles i.e *χ* and Θ. We obtained a converged 2D PMF along *χ* and Θ torsion angles using meta-eABF simulations within 200 ns for both the DNA and RNA system. Convergence was monitored by plotting the root mean squared deviation (RMSD) of the PMF at 200 ns from that at earlier times (Figure S1 in Supporting Information). The presence of HG base pairing in DNA is reflected by the clear deep minimum at *χ* ~ 50° and Θ ~ 0°. The Watson-Crick base pair is located in the deep stretched well between −180° and −50° of the glycosidic angle *χ* for Θ close to 0°. Meanwhile, for the RNA the WC base pairing was observed at *χ* ≈ −160°. Additionally, there is a shallow minima at *χ* ≈ 30° in RNA, close to that of HG base bair in DNA (Fig. 1). Careful examination of the trajectory revealed a Hoogsteen like structure in RNA with only one hydrogen bonding between NH_2_ of adenine and carbonyl oxygen of thymine (distance ≤ 3.0 Å and angle ≤ 30°) (Fig. 1). This HG like structure is energetically unstable by ~ 2 kcal/mol compared to the HG base pair of the DNA. This structure has previously been observed by Rangadurai et. al. by computer simulation (64) and is referred to as RNA HG base pair in the rest of the paper. RNA also showed a higher free energy barrier between the HG and WC form.

**Figure 1:**
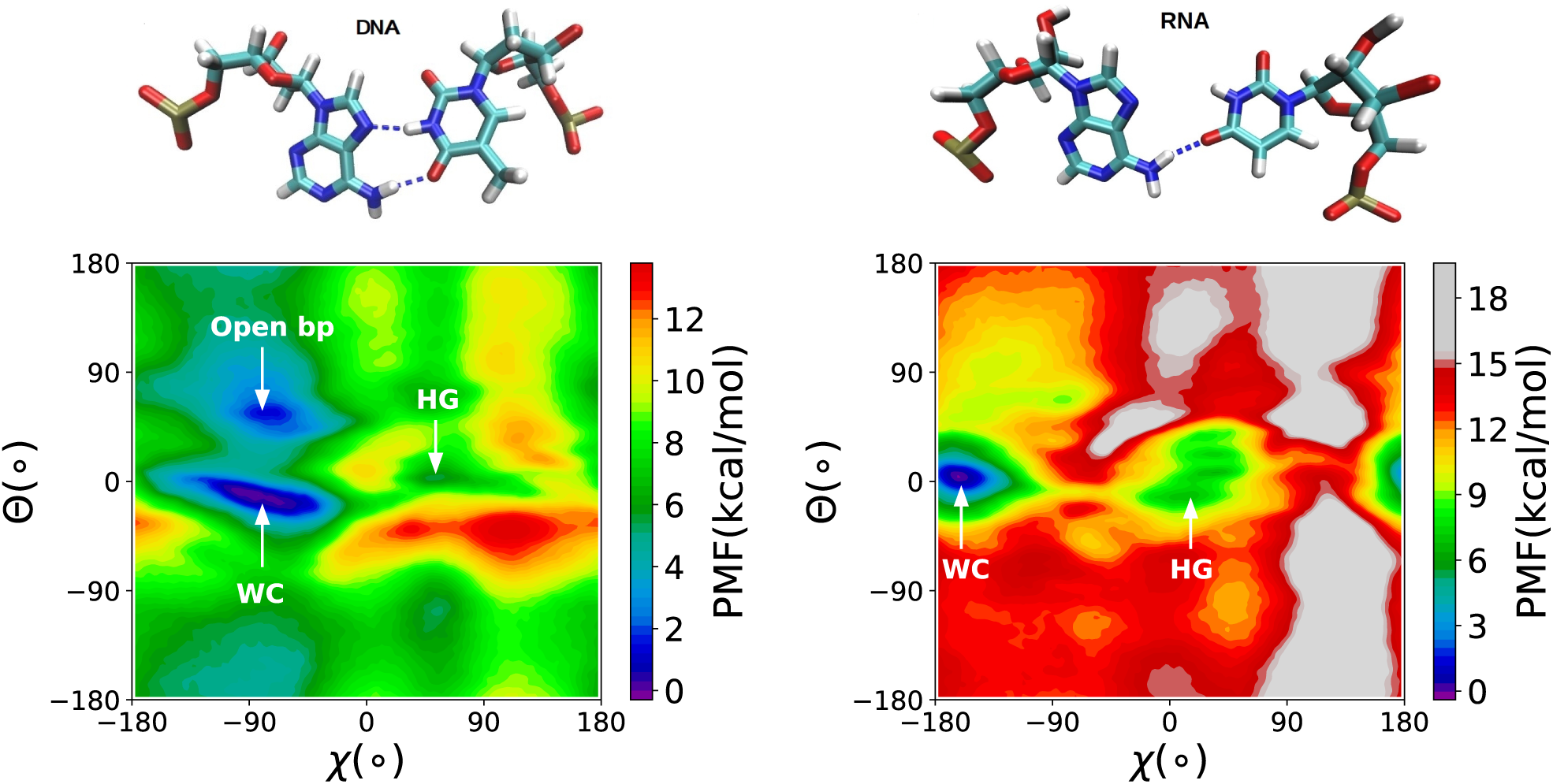
Upper panel: The structures of Hoogsteen and Hoogsteen like base pair in A6-DNA and A6-RNA. Lower panel: 2D PMF of the A16-T9 base pair of the A6-DNA and A16-U9 base pair in A6-RNA computed using meta-eABF method.

The experimental observation of RNA melting by methylation of adenine can be attributed to the relative instability of the RNA HG base pair although we did not observe any RNA helix melting in our simulations. The simulated free energy differences and barrier heights are not directly comparable with the experimental results which are estimated based on simple assumption of Boltzmann probability and transition state theory (TST). Still, we obtained the relative free energy of the HG base pair with respect to WC (∆*G*_*WC* → *HG*_) to be 4.5 kcal/mol which is within 1 kcal/mol agreement with NMR experiments and microsecond timescale umbrella sampling and meta-dynamics simulations (23, 25, 26). The positions of the WC and HG minima in *χ* − Θ space also agree with earlier computational results (25, 26). The absence of the N7(A)-N3(U) hydrogen bond can be the possible reason which makes RNA HG unstable (64) and elusive in experiments.

The difference of relative free energy of the HG/HG-like base pair between DNA and RNA is more apparent in the *θ* integrated 1D PMF along as a function of *χ* angle (Figure 2). The PMF along the base flip-out angle Θ shows a second deep minimum at Θ ~ 60° connected to the WC base paired state with a small barrier in DNA (Figure 2). It is in agreement with the work of Lavery and coworkers (65) who obtained a similar free energy minimum for adenine base opening through major groove via what they called a “saloon door” mechanism.

**Figure 2:**
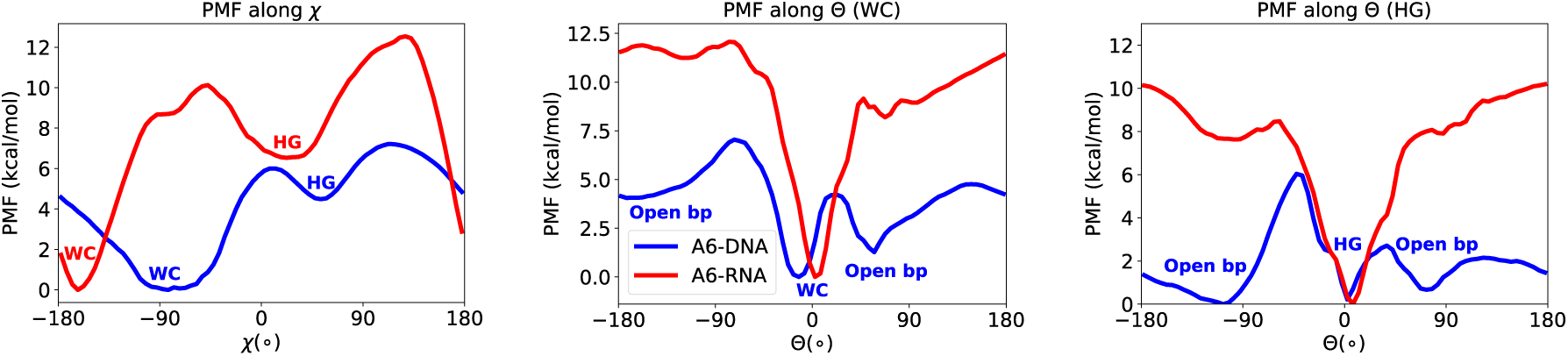
One dimensional PMF’s along glycosidic angle *χ* and pseudo dihedral angle Θ angle for WC and HG base pairing for both A6-DNA and A6-RNA computed using Equation 1.

NMR experiments also observed spontaneous base pair opening in DNA duplexes (66) at a much faster rate than the formation of HG base pairing (2). Such consistencies with previous results bolster our claim of capturing the essential physics of DNA base pair dynamics from relatively short meta-eABF simulation. Two minima corresponding to open base pair conformations are present, one towards the major and the other towards the minor groove side of the HG base pair. However, the free energy barrier of base pair opening through the minor groove 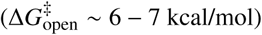 is much higher than through the major groove 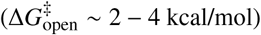. This is in agreement with the results of Guidice et al. (65) and our own results in Nikolova et al. (2), which show that the base pair opening and flipping motions take place through the major groove (positive Θ) of DNA predominantly. We do not see any such low energy minima of open base pair conformations for RNA (Fig. 2).

A clearer indication of base pair configuration can be obtained by looking at the the hydrogen bond donor-acceptor distances. In WC base pair, hydrogen bond forms between the N1 of A and N3 of U/T base, while for HG base pairing, N7 of A participates in hydrogen bonding instead of N1. We have plotted the N1-N3 and N7-N3 bond distances for a small stretch of the meta-eABF trajectory for both DNA and RNA (Fig. 3). Shorter distances (~ 3.5 Å) between N1-N3 or N7-N3 atoms correspond very well with the real WC and HG base paired conformation respectively. The agreement of *χ* and Θ angles is not as good as the distances between N atoms and there are certain structures with HG or WC specific *χ* and Θ values, although they are not base paired structures. This can be due to the fact that this angles are measured with respect to the sugar or the neighbouring base pairs whose structures also fluctuate with time. It also indicates that in order to get a correct picture of the transition between WC and HG base pair we should take hydrogen bond donor acceptor distances into account. Figure 3 also shows the variation of the helix diameter measured by the distance between backbone C1’ atoms of the participating nucleotide. The distance is about 10 Å for WC base paired forms in DNA and RNA. For HG base pairing in DNA, the helix diameter shrinks to ~ 8.5 Å in agreement with previous results (58). Interestingly for RNA HG like structure this increases to ~ 12 Å and the N1-N7 distance does not shrink as much as it is in DNA. These are the structural factors which prevent the formation of the second hydrogen bond in the RNA HG. The effects can be a result of the larger diameter of the A form helix, present in RNA, than the naturally occurring B form in DNA. So the free energy compensation for the shrinkage of radius in RNA helix is large. This has been attributed to be the reason for the absence of HG in RNA in a recent experimental and computational study (64).

**Figure 3:**
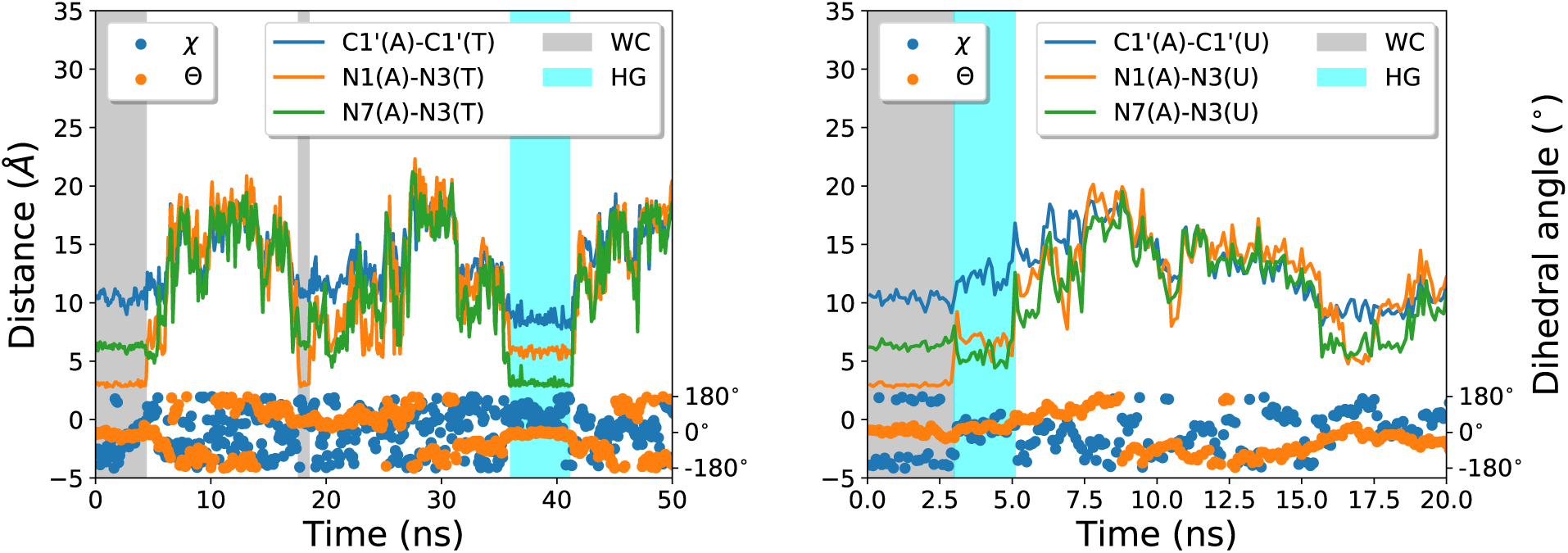
Formation of Hoogsteen base pairing in the meta-eABF simulation. The hydrogen bond donor-acceptor distance, the distance between *C*1′ atoms of the backbone corresponding to the adenine and thymine bases and the *χ* and Θ torsion angles are shown for the first few frames of the trajectory for DNA (left) and RNA (right). The regions of WC and HG base pairs are indicated by manual inspection of the trajectory frames. Similar behaviour is also observed throughout the 200 ns trajectory but only a small portion is shown for clarity.

### Markov State Modeling

#### TICA analysis and reaction coordinate

Given that the traditional torsion angles *χ* and Θ remain inadequate to appropriately describe the conformational transition between WC and HG base pairing forms, a multidimensional representation involving other structural features becomes necessary. Time lagged independent component analysis (TICA) can be used to identify the slowest degrees of freedom in a complex conformational transition as a function of internal coordinates of the system (43, 44). TICA analysis of our unbiased trajectory data predicted the slowest independent components (IC) in DNA base pair motion are strongly correlated with the hydrogen bond donor acceptor distances (Figure 4). The third H-bond distance between (T/U) O4 - (A) N6, which is common in both WC and HG, also showed high correlation indicating that both hydrogen bonds get broken during the slowest motions of the base pair.

**Figure 4:**
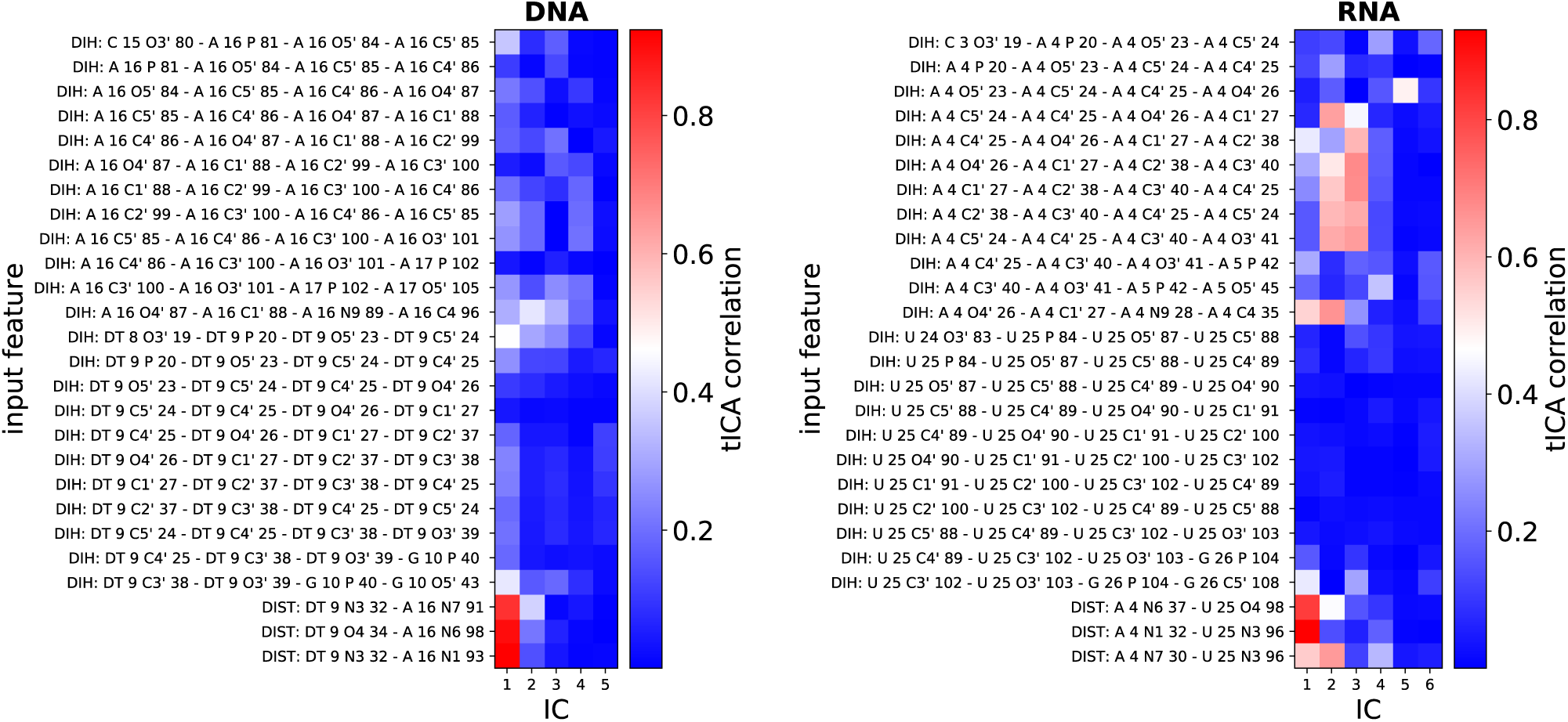
Absolute value of the correlation coefficient of the independent components (IC) obtained from TICA analysis with the input features i.e dihedral angles and possible hydrogen bond donor-acceptor distances for DNA and RNA.

Among the dihedral angle based features, the phosphodiester bond torsion angles have highest correlation with IC 1 in DNA, possibly indicating slow base flip-out motions. Conversely none of the IC’s for RNA has significant correlation with the torsional angle of phosphodiester bond. In DNA the IC 2 and IC 3 shows high correlation with the glycosidic angle *χ* suggesting that IC 2 is a better coordinate than IC 1 for resolving the WC to HG transition. Meanwhile, in RNA both IC 1 and IC 2 are strongly correlated with the *χ* angle. Also IC 2 and 3 has significant contribution from the sugar pucker angles in the ribose sugar of the adenine nucleotide. None of the torsional angles in the uracil nucleotide has any contribution in the slow motions captured by our TICA analysis.

#### MSM structural network, free energy and kinetics

Markov State Model (MSM) (40) decomposes unbiased molecular dynamics trajectories into clusters based on structural criteria and calculates transition probabilities between them. From this transition probability data, free energy landscape, conformational kinetics and mechanistic pathways can be elucidated (67). We constructed an MSM based on structural features using the internal coordinates (IC) obtained from TICA. The MSM was decomposed into 5 and 4 meta-stable states respectively for DNA and RNA to get an intuitive understanding of the pathway of transition. They were identified as WC, HG or intermediate (I) states by inspection of sample structures. The choice of the number of meta-stable states were optimum for differentiating between recognizable structural forms (i.e. WC, HG, base pair open etc). To analyze these states we define a base flip out angle for thymine/uracil (*η*) analogous to the Θ angle for adenine. The meta-stable states were then projected onto both *χ* − Θ and Θ − *η* space to understand their correspondence with the free energy landscape obtained from meta-eABF simulation (Figure S9, S10 and S11 in Supporting Information). For DNA, the meta-stable states overlap each other significantly in the *χ* − Θ space and many open base conformations coincide with both the WC and HG states. This result is confirmed by Θ − *η* plots where a base paired state is indicated by a cluster close to the origin. If either of the base is in extra-helical conformation the corresponding pseudo-dihedral angle will be away from zero (Figure S10 and S11 in Supporting Information). The I1 has at least one of the base is in open conformation while I3 has both of them open. I2 primarily has the adenine inside the helix (Θ ~ 0) but the thymine is outside (Figure 6). But, both I1 and I2 coincide with base paired state in the *χ* − Θ space leading to over-counting the population of WC and HG. These results indicate that conventional dihedral angle based representation fails to capture the true story behind the formation of HG base pairing. This limitation is, most probably, a result of not taking the conformation of the thymine base into account.

Unlike DNA, none of the meta-stable states of RNA showed extra-helical conformation. Although in intermediate I the inter base hydrogen bonds are broken, there is a new hydrogen bond between the uracil carbonyl group oxygen and the *C*2′ -OH group in the ribose sugar of the adenine nucleotide (Figure 7). This intermediate is further stabilized by another hydrogen bond forming between adenine (N3) and the -OH group of phosphodiester bond in the backbone of uracil nucleotide. They prevent the adenine to completely flip out of the helix. Consequently the extra-helical conformations are unfavourable in RNA A-T base pair because of the free energy cost of breaking these additional hydrogen bonds.

From the Markov state model, we computed the free energy landscape for the A-T/U base pair conformation as a function of the two slowest degrees of freedom obtained from TICA analysis (Figure 5). Comparing the results with the meta-stable state distribution we can get an idea of the configurations represented by different minima in the free energy surface. Clear deep minima for both WC and HG base pairing were obtained in DNA while, in RNA, apart from a HG like state we observed two different WC minima (Figure 5). One of them has the ribose sugar in *C*3′-endo conformation (WC 1) and the other in *C*2′-endo conformation (WC 2) (Figure 7). As dictated from the involvement of sugar pucker angles in the first TICA component, the transition from *C*3′-endo to *C*2′-endo conformation is one of the slowest motion captured in our base pair dynamics model of RNA. The relative free energy of HG (∆*G*_*WC* → *HG*_) obtained from MSM agrees with the results of meta-eABF and with previous experimental and computational studies. However, this agreement can be a coincidence considering that the WC and HG minima in the *χ* − Θ based PMF have contribution from non base paired intermediates.

**Figure 5:**
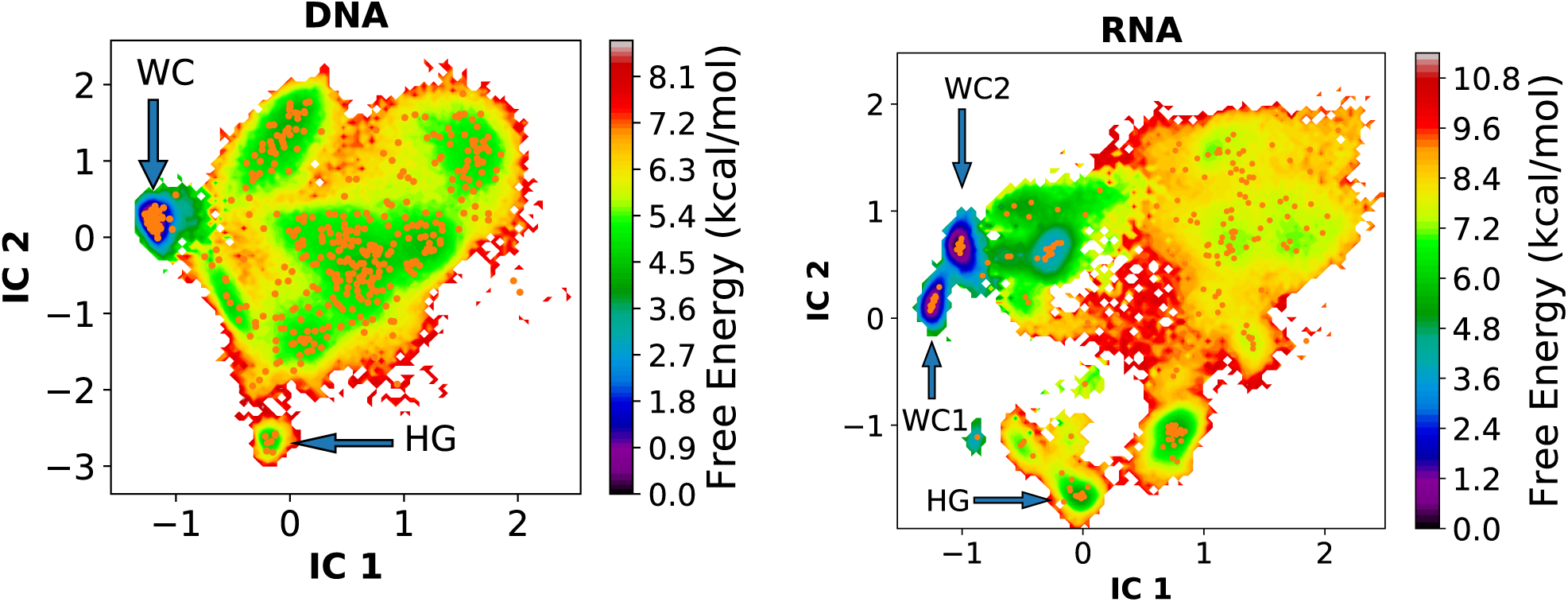
Free energy profile of A-T base pair conformations for both DNA and RNA as a function of the two slowest independent components (IC) obtained from TICA based MSM.

We utilized transition path theory (TPT) (62, 63) to obtain the kinetics between the clusters. The kinetic information was projected onto the coarse grained space to generate a flux network between various meta-stable states (Figure 6 and 7). The mean first passage time of transition from WC to HG state in DNA was obtained to be 1.8 ms, which is in great agreement with the experimental lifetimes of HG base pair (0.2-2.5 ms) (68) but an order of magnitude less than the experimental first passage times (~ 50 ms) (2). Our result also agrees quite well with the recently reported rate constants of transition between WC and HG base pairing using TPS (27). A simple Arrhenius theory based estimation yields an activation free energy of ~ 14 kcal mol^−1^ (Table 1) which matches well with such estimates made from NMR experiments (~ 16 kcal mol^−1^) (2). The majority of the flux in the DNA system is along the following two paths: *HG → I*3 → *WC* (87.2%) and *HG → I*3 → *I*2 → *WC* (8.8%). It indicates that the most probable intermediate between WC and HG is the state with both base pair flipped outside the helix. It also provides qualitative evidence that the transition between WC and HG base pair happens through concerted outwards movement of the bases followed by coming back inside the helix, in agreement with earlier propositions (2, 27).

**Table 1:** Mean first passage times (MFPT) and barrier heights for transition between WC and HG base pairing forms predicted from MSM for both A6-DNA and A6-RNA.

|  | WC→HG |  | HG→WC |  |
| --- | --- | --- | --- | --- |
| | MFPT (ms) | Barrier height (kcal/mol) | MFPT (μs) | Barrier height (kcal/mol) |
| DNA | 1.8±0.4 | 13.89±0.14 | 0.85±0.07 | 9.28±0.05 |
| RNA(WC1) | 30.2±10.8 | 15.57±0.22 | 10.7±3.85 | 10.76±0.23 |
| RNA(WC2) | 32.0±11.7 | 15.60±0.23 | 567±70 | 13.18±0.07 |

**Figure 6:**
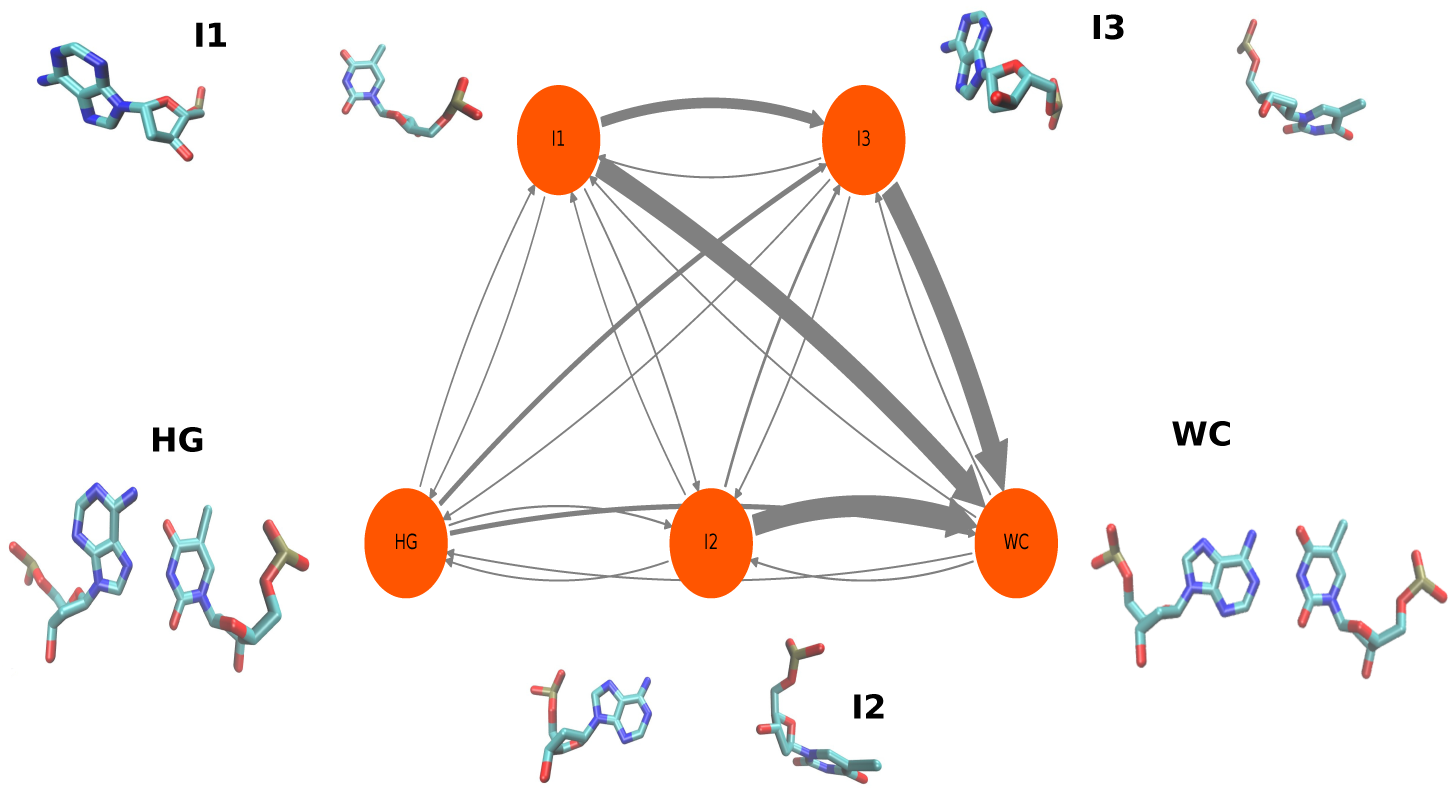
A network of fluxes through the five meta-stable states obtained from MSM for A6-DNA system

**Figure 7:**
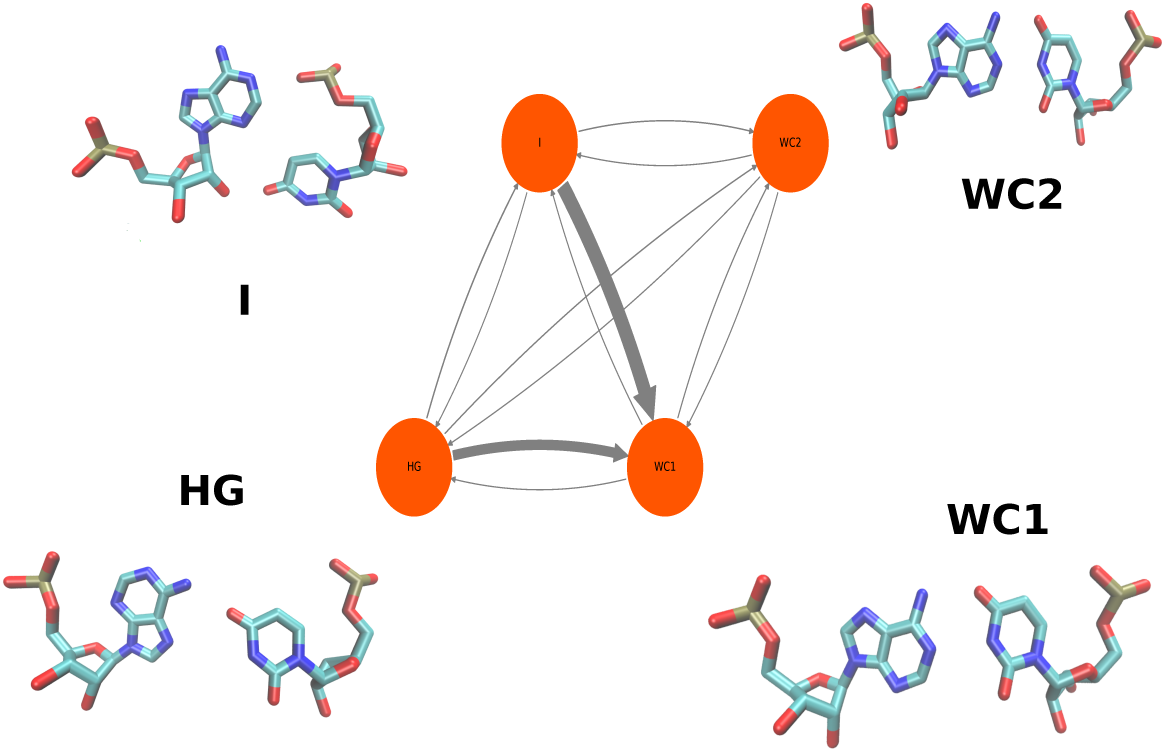
Same as Figure 6 but for A6-RNA system.

Our model predicts the transition timescale to HG base pair from WC1 to be 30.2 ms and from WC2 to be 32.0 ms for RNA system (Table 1), although there is no experimental data available to compare. The timescales of the NMR *R*_1*ρ*_ relaxation experiments are a few tens of milliseconds (69). So the transition timescale from WC to RNA HG is close to the maximum limit of this experiments and consequently, it suggests the reason why a HG-like base pair has not been experimentally detected in the A6-RNA hairpin. The major pathways of probability flux are the following: *HG → I → WC*1 → *WC*2 (84%) and *HG → WC*1 → *WC*2 (16%). None of the intermediates assume open base pair conformation which is evident from the Figure 7 and from the Θ − *η* distribution of the meta-stable states. The transition between WC and HG-like base pairing in RNA takes place through an intra-helical pathway. Our 2D PMF from meta-eABF simulation also supports the claim as the free energy cost of A-U base pair opening RNA is observed to be much higher than that of A-T base pair in DNA (Figure 2).

## CONCLUSION

In this work, we combined the meta-eABF enhanced sampling technique with Markov state modeling to obtain mechanistic insights into the conformational switching between WC and HG base pairing in DNA and RNA. We have constructed and compared the 2D PMF along the glycosidic and base flip out angles, which showed presence of HG or HG like conformation in both DNA and RNA. Markov state modeling produced a free energy landscape along the two slowest degrees of freedom with distinct free energy minima for WC and HG base pair for DNA. It predicted the relative free energy of HG base pair within ~1 kcal/mol accuracy of previous experimental and computational results. We also observed single hydrogen bonded and relatively unstable HG-like base pairing in RNA. The absence of the second hydrogen bond can be attributed to the larger diameter of A-form helix in RNA which does not allow for sufficient shrinkage of the hydrogen bond donor-acceptor distance. Monitoring the distance between the backbone *C*1′ atoms in adenine and thymine/uracil nucleotide, from the biased trajectory, substantiated our conclusion about insufficient shrinkage of RNA helix diameter in HG like base pairing.

Our inference is in agreement with recent arguments made by Rangadurai et el. who showed A form of helix needs to perform a significant sugar-backbone rearrangement to avoid steric clash in the base in *syn* form present in HG (64).

A closer look at the biased trajectory revealed that the WC and HG state definitions correspond better with the hydrogen bond donor-acceptor distance than with the dihedral angles, traditionally chosen as collective variables. TICA analysis of multiple unbiased trajectories and dihedral angle distribution of the meta-stable states from Markov state model (MSM) agree with this conclusion. The slowest dynamics of the system involves forming and breaking of the hydrogen bonds, and also conformational changes of the thymine/uracil base. The relative free energy of the RNA HG with respect to the DNA HG is ~2 kcal/mol which is roughly the energy of one hydrogen bond in nucleic acid base pairing (70). Although observed in protein and drug bound DNA and synthetic DNA constructs, Hoogsteen base pair in RNA remained elusive in experimental studies. The ~6.5 kcal/mol free energy difference with respect to WC base pairing form makes RNA HG extremely rare (< 0.002% according to Boltzmann distribution arguments) and unlikely to be observed with present experimental capabilities. The observation of RNA HG in our study is of particular significance because our enhanced sampling scheme was not biased towards any specific structure (for example, as typically done in targeted MD (71) or other biased methods. It shows that there is an inherent stability of the HG like structure in RNA and is not a simulation artifact.

MSM constructed from TICA coordinates predicted free energy and kinetics of transition between WC to HG base pair in A6-DNA within reasonable agreement with previous theoretical (25–27) and experimental results (2). But we also predicted that the timescales for such transition in RNA is more than 30 ms. Considering that the MSM timescale for WC to HG transition in DNA is one order of magnitude less than the experimental data, the true first passage time for RNA might go well beyond the time range of NMR *R*_1*ρ*_ relaxation experiment. This provides a parallel kinetic rationale for why HG like base pairing is not observed in experimental studies. A kinetic network was constructed between a handful of structurally distinguishable meta-stable states obtained by decomposing the MSMs. Most probable transition pathway, traced down from the networks, indicate that the conformational switching between Watson Crick and Hoogsteen base pair happens through an extra-helical mechanism for A6-DNA while in RNA hairpin intra-helical mechanism is dominant. This result is explained by the much higher free energy cost of base pair opening in RNA compared to DNA obtained from the PMF using meta-eABF simulation. The analysis of pathway also resulted in the observation of novel intermediate structure with unusual backbone-base hydrogen bonding in A6-RNA.

Application of meta-eABF method resulted in accurate PMF in two orders of magnitude less computational effort for HG base pairing. This method shows promise to be used extensively over a range of protein-DNA and drug-DNA complexes with HG base pairing, and make comparison between ∆*G*_WC→HG_ free energies possible within reasonable computing time. Pathways and timescales of HG pairing in such systems can be understood from Markov state models and multidimensional path searching methods like TPSS (72), String methods (73–75) and other similar techniques (76, 77). Taking additional degrees of freedom like H-bond donor-acceptor distances, sugar pucker angles, and phopho-diester bond torsion angle into consideration, will provide better understanding of the molecular processes involved in HG base pair formation.

We hope our work will motivate careful experiments in both short and long timescales for the detection of HG base pair and novel short-lived intermediates in duplexes of RNA hairpins. This work, consequently, marks the beginning of a large scale exploration of Hoogsteen base pairing thermodynamics and kinetics in various biologically relevant nucleic acid systems, through computational and subsequent experimental methods, to gain detailed mechanistic understanding of the role of non-canonical base pairing in nucleic acid function and recognition.

## AUTHOR CONTRIBUTIONS

D.R. and I.A. designed research; D.R. performed simulation and analyzed data with inputs from I.A.; and D.R. and I.A. wrote the paper.

## ACKNOWLEDGMENTS

This work has been supported in part by funds from the National Institutes of Health, grant RO1GM089846-08. The authors thank James McSally for help in building the initial structures of DNA and RNA. The authors thank Christophe Chipot and Victoria Lim for helping set up meta-eABF calculation and Nicolae-Viorel Buchete, David Mobley, Sam Gill and Chris Zhang for valuable suggestions regarding Markov State Modeling. The authors thank San Diego Supercomputer Center (SDSC) for computational resources.

## Supporting information

### Theoretical Details of meta-eABF

In the eABF method by Lesage et al. [1], a fictitious particle is attached to the collective variable by a strong force constant. An external force, equal and opposite to the potential energy gradient, is applied to the fictitious particle, which forces the dynamics against the potential energy gradient. To give a brief overview, the extended potential energy *V*^ext^(*q, λ*) in presence of the harmonic coupling with fictitious coordinate *λ* is given by

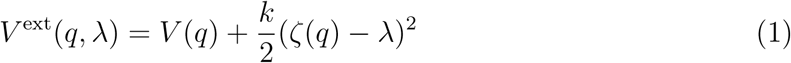

where *q* is the system cartesian coordinates and *ζ*(*q*) is the order parameter or transition coordinate as a function of system coordinates whereas *k* is a strong coupling constant between the reaction coordinate and the fictitious coordinate. An external force *F*^*bias*^ is the applied on the coupled coordinates as

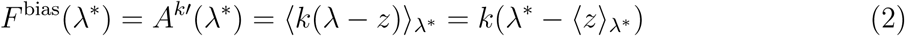

where angular brackets denote conditioned average constrained by *λ* = *λ*^∗^ and *A*^*k*^(*λ*) is the PMF along coordinate *λ* defined as

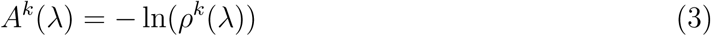

So the time dependent biased potential as function of *q* and *λ* converges to

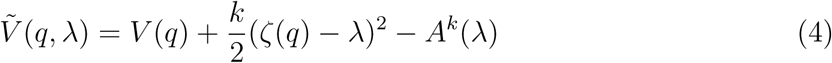

leading to a the Boltzmann probability along *z* = *ζ*(*q*)

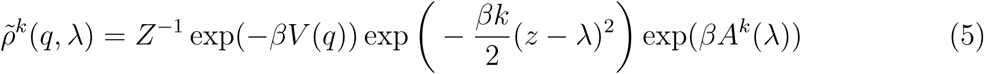

where 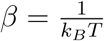 and *Z* is canonical partition function. Now identifying that *ρ*(*z*) *∝* exp(*−βV* (*q*)) and 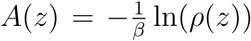 and also using the Bayes theorem for the conditional probability 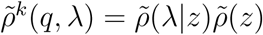 it is concluded that the first derivative of free energy along the coordinate z is given by

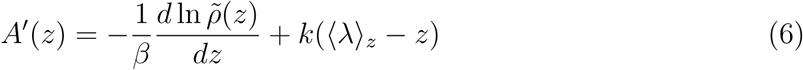

This is known as corrected z-averaged restraint (CZAR) estimator expression which is integrated numerically to obtain the PMF.

The eABF method alone was not sufficient to accelerate the dynamics enough to sample the configuration space within reasonable computational cost. So, metadymanics (MtD) [2] was performed along with eABF to gain further acceleration by simultaneously depositing Gaussian bias on the reaction coordinate. So the net biasing force is given by the following expression: [3]

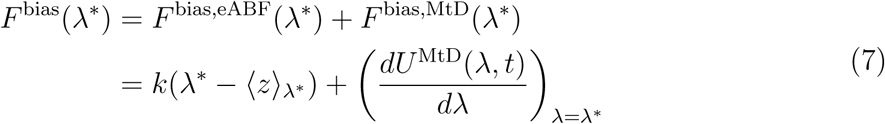

where *U*^MtD^(*λ, t*) is the time dependent biasing potential deposited by MtD run. The energy gradient obtained using this process has contribution from both MtD and eABF. The potential energy gradient and the number of counts in each collective variable bins were stored and integrated using abf_integrate package with CZAR estimator to calculate the potential mean force (PMF). Although the indiviudual contribution from MtD and eABF to the PMF may change with time but the total PMF remains constant once the simulation has converged [3].

---

**Figure S1:**
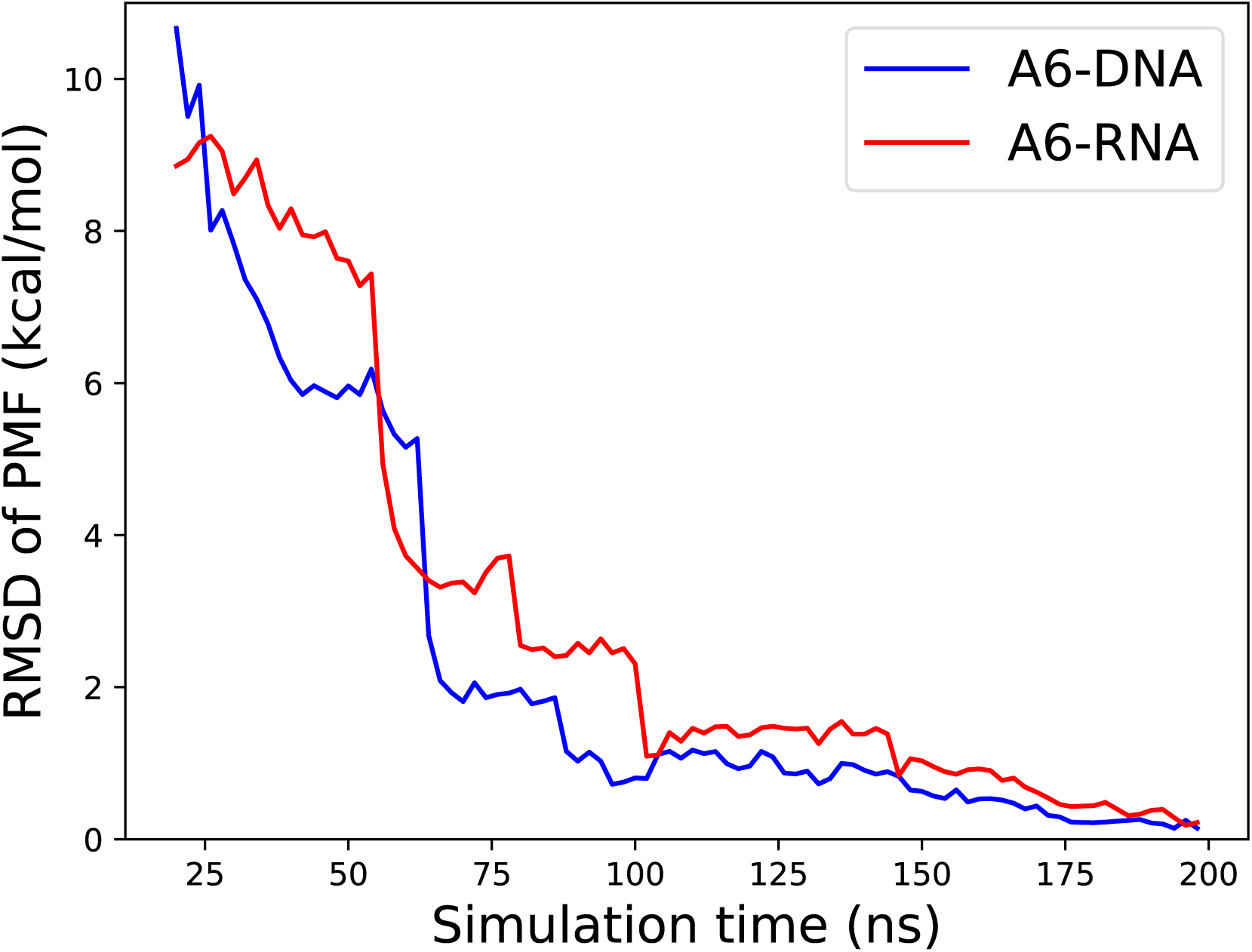
Convergence of the 2D PMF calculated from meta-eABF simulation for DNA and RNA system.

**Figure S2:**
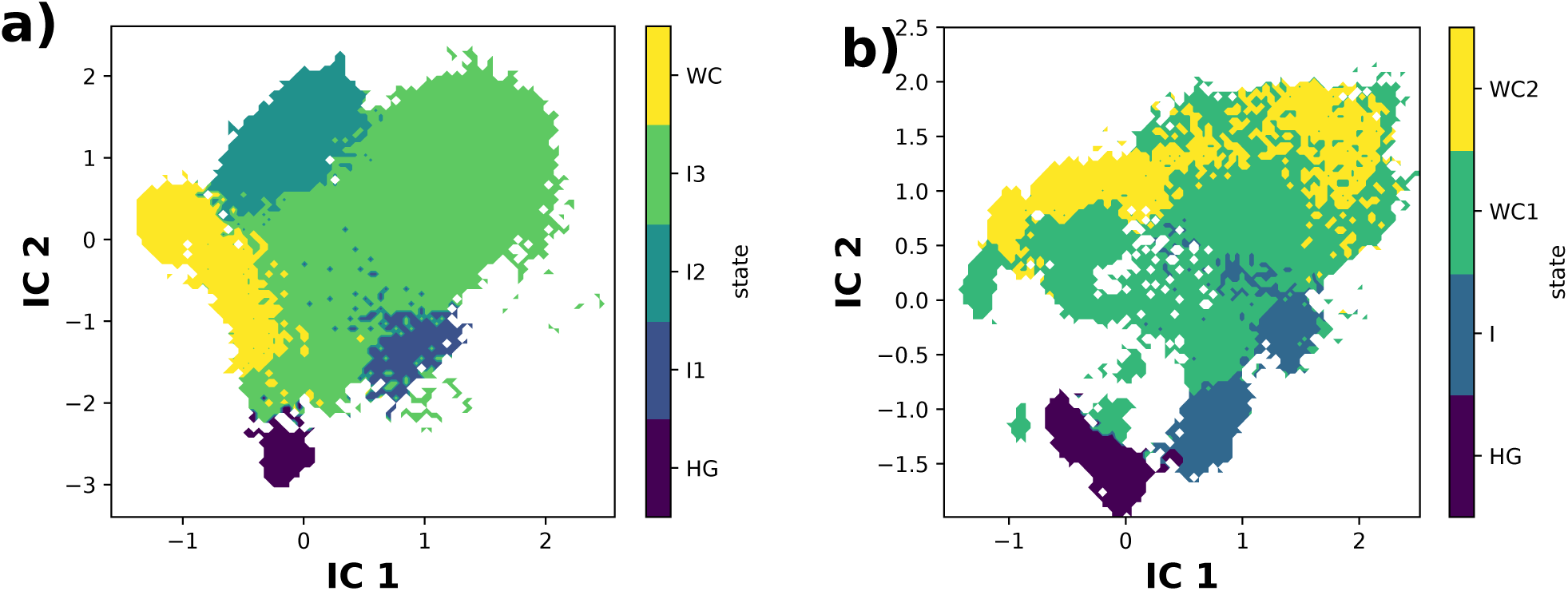
Coarse grained meta-stable states as a function of the two slowest independent components (IC) for a) A6-DNA and b) A6-RNA

**Figure S3:**
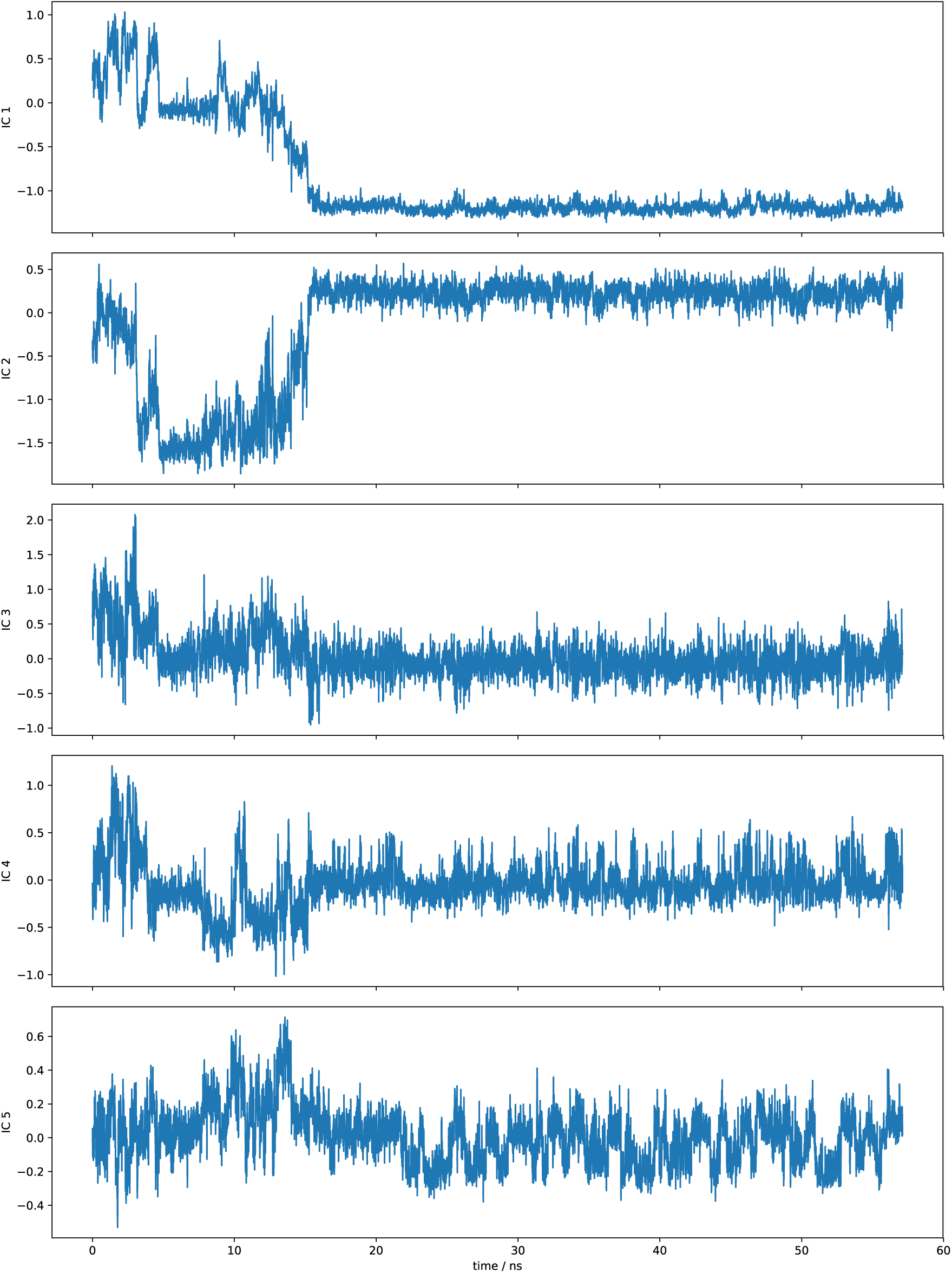
Fluctuation of each IC between metastable states for A6-DNA

**Figure S4:**
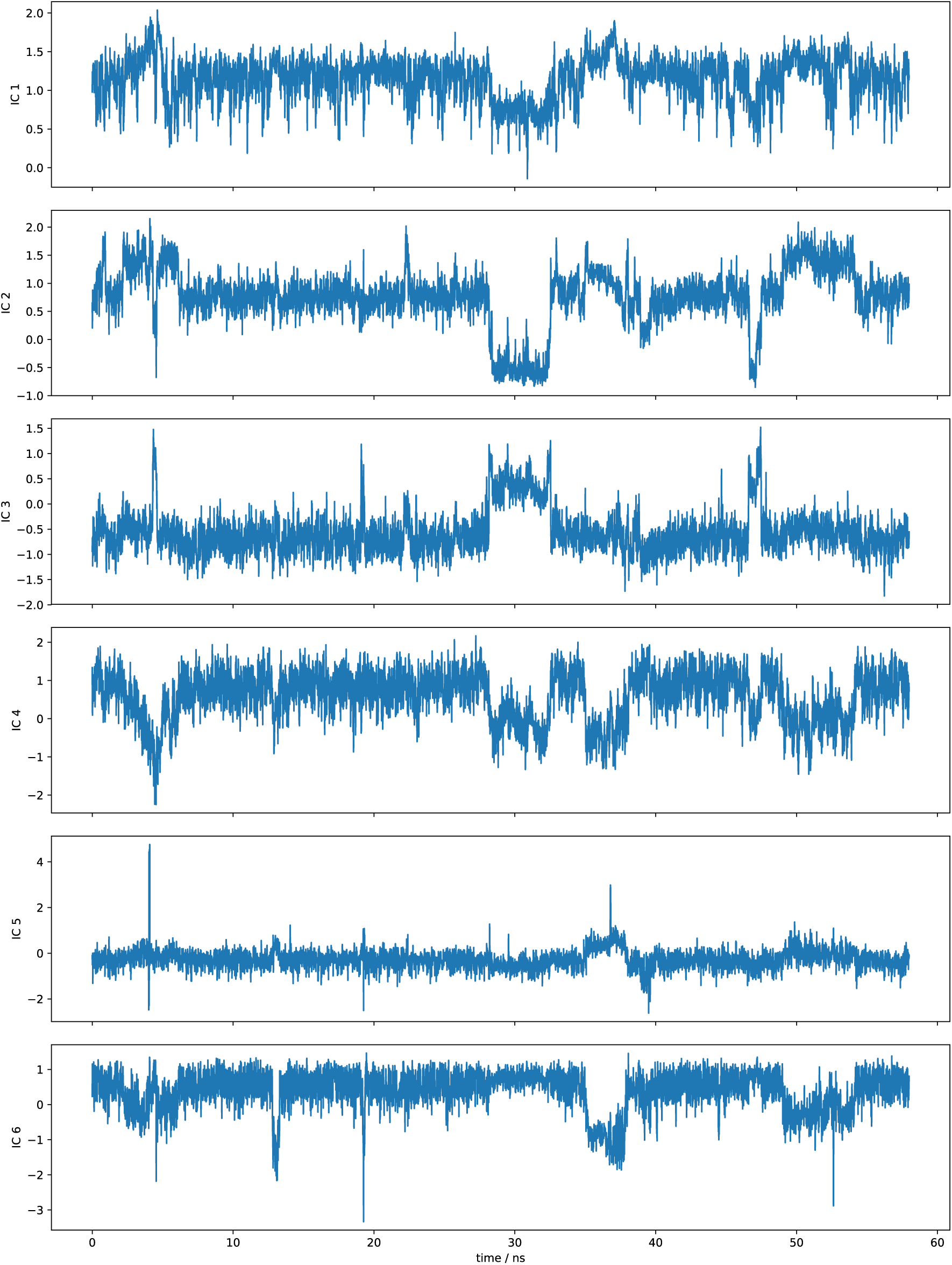
Same as Figure S3 but for A6-RNA

**Figure S5:**
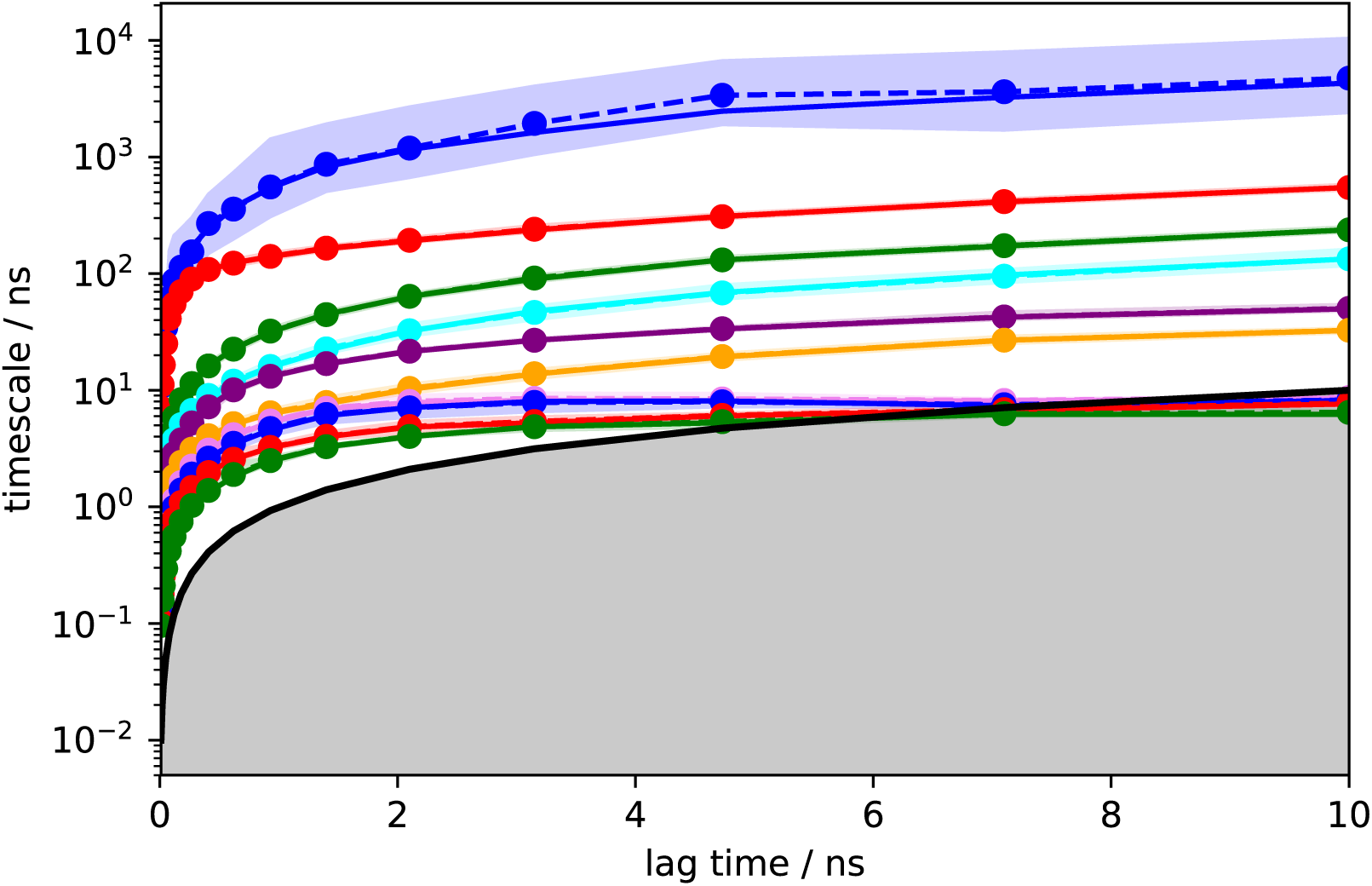
Implied timescale convergence of the 10 slowest eigenvalues of the Markov state model constructed for A6-DNA. Solid lines indicate average data and errors are indicated as shaded area around the curves.

**Figure S6:**
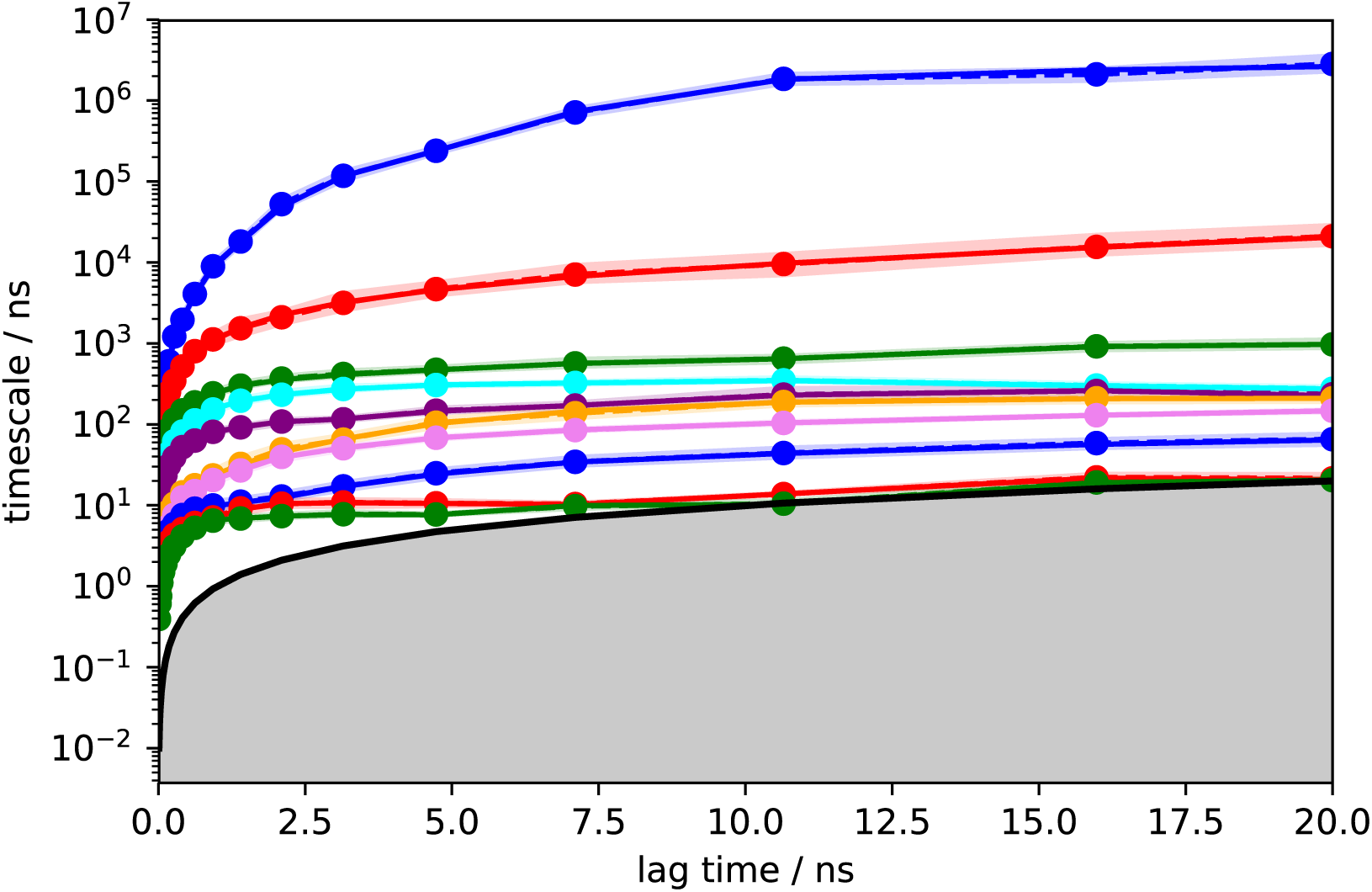
Same as Figure S5 but for A6-RNA.

**Figure S7:**
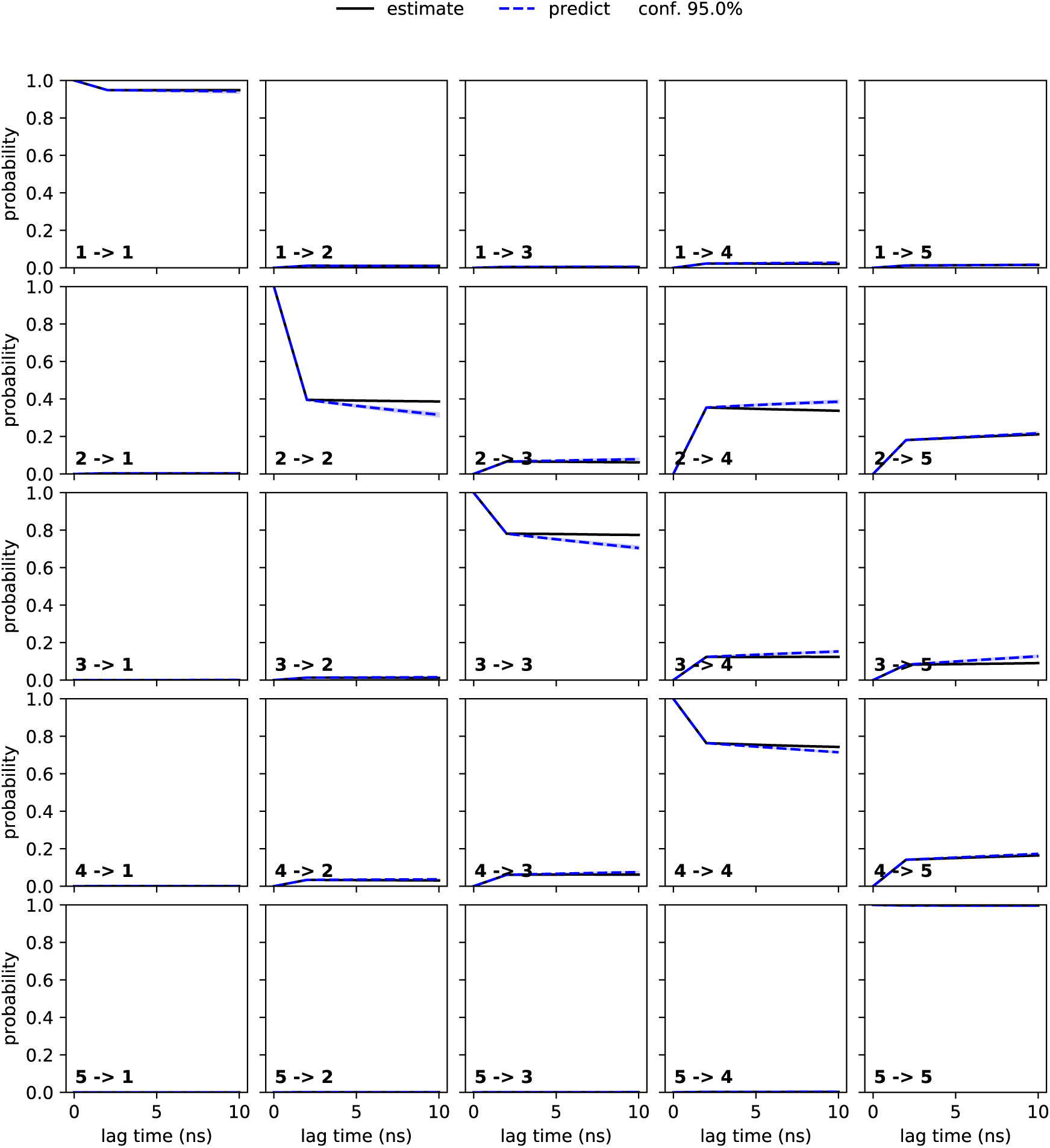
Results of Chapman Kolmogorov Test for the Markov model constructed for A6-DNA.

**Figure S8:**
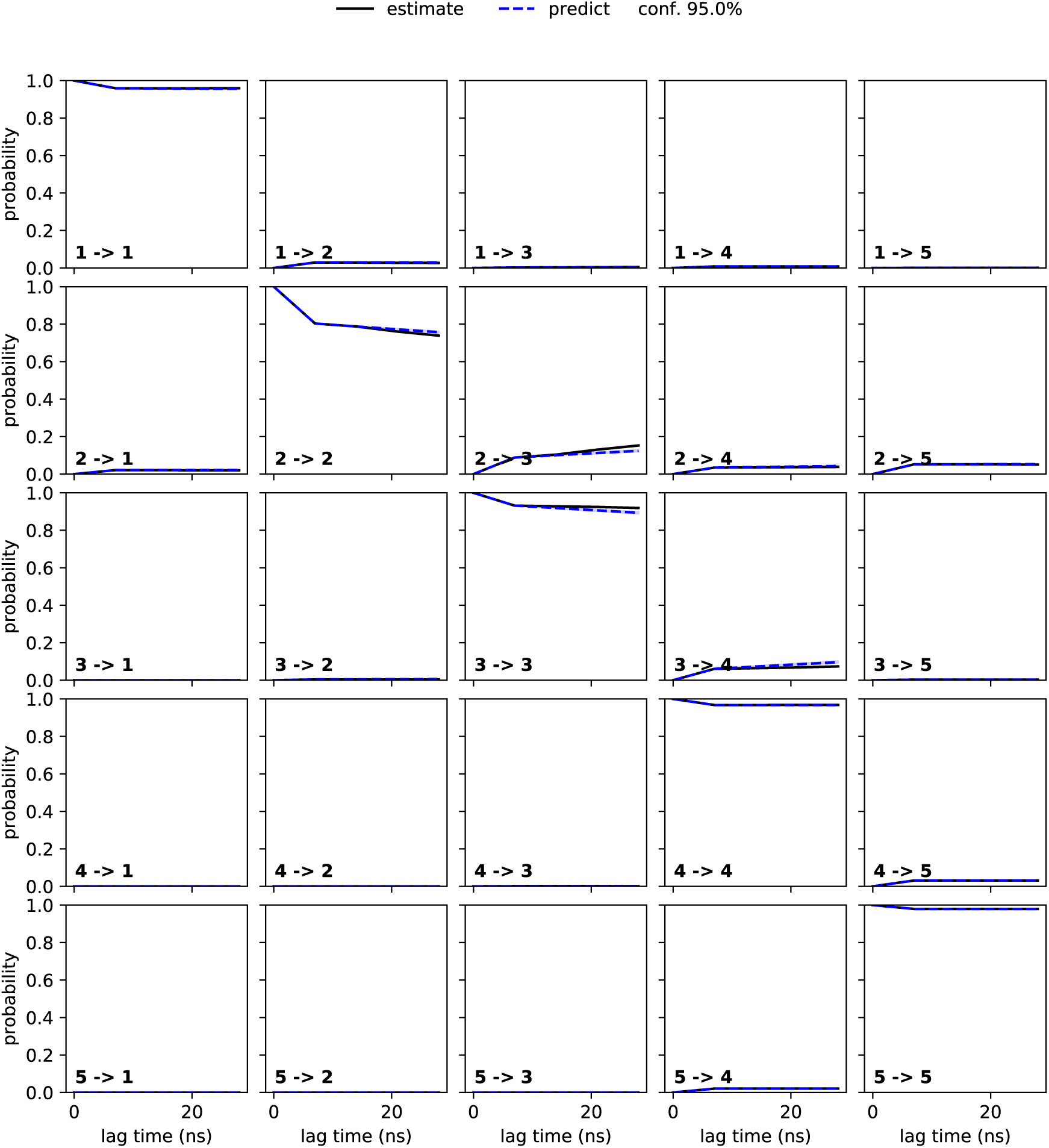
Same as Figure S7 but for A6-RNA.

**Figure S9:**
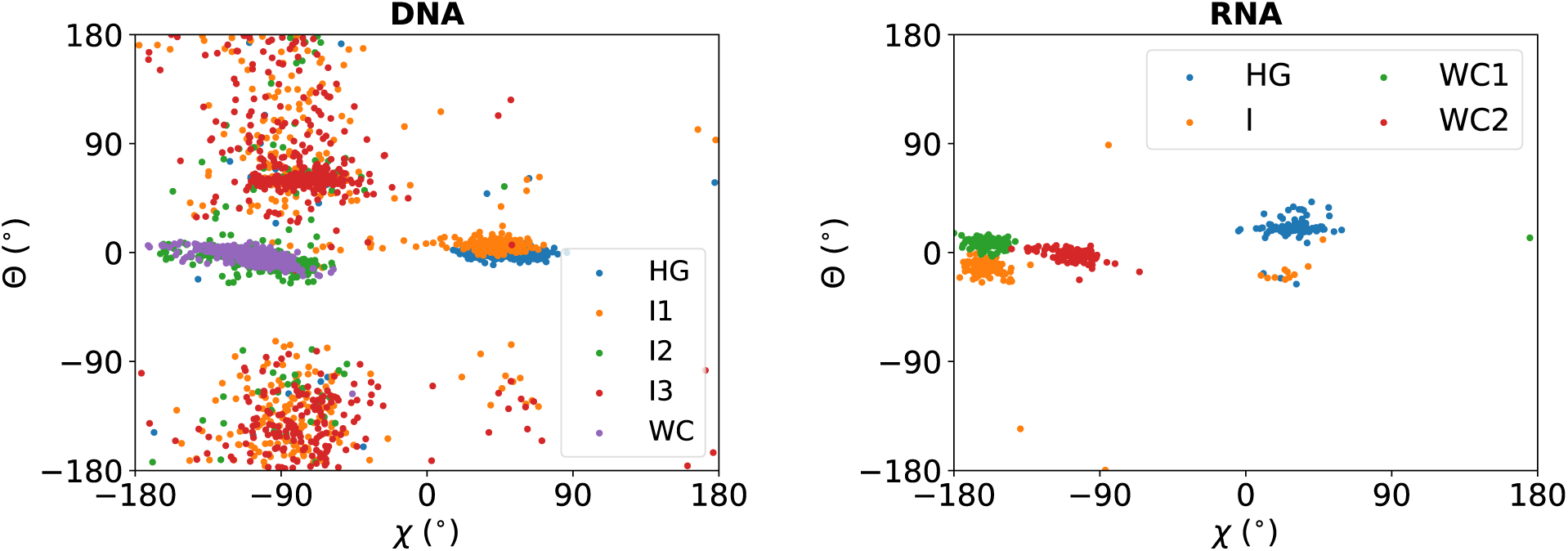
Distribution of glycosidic angle **χ** and pseudo dihedral angle Θ for the meta-stable states obtained from MSM of A6-DNA (left) and A6-RNA (right)

**Figure S10:**
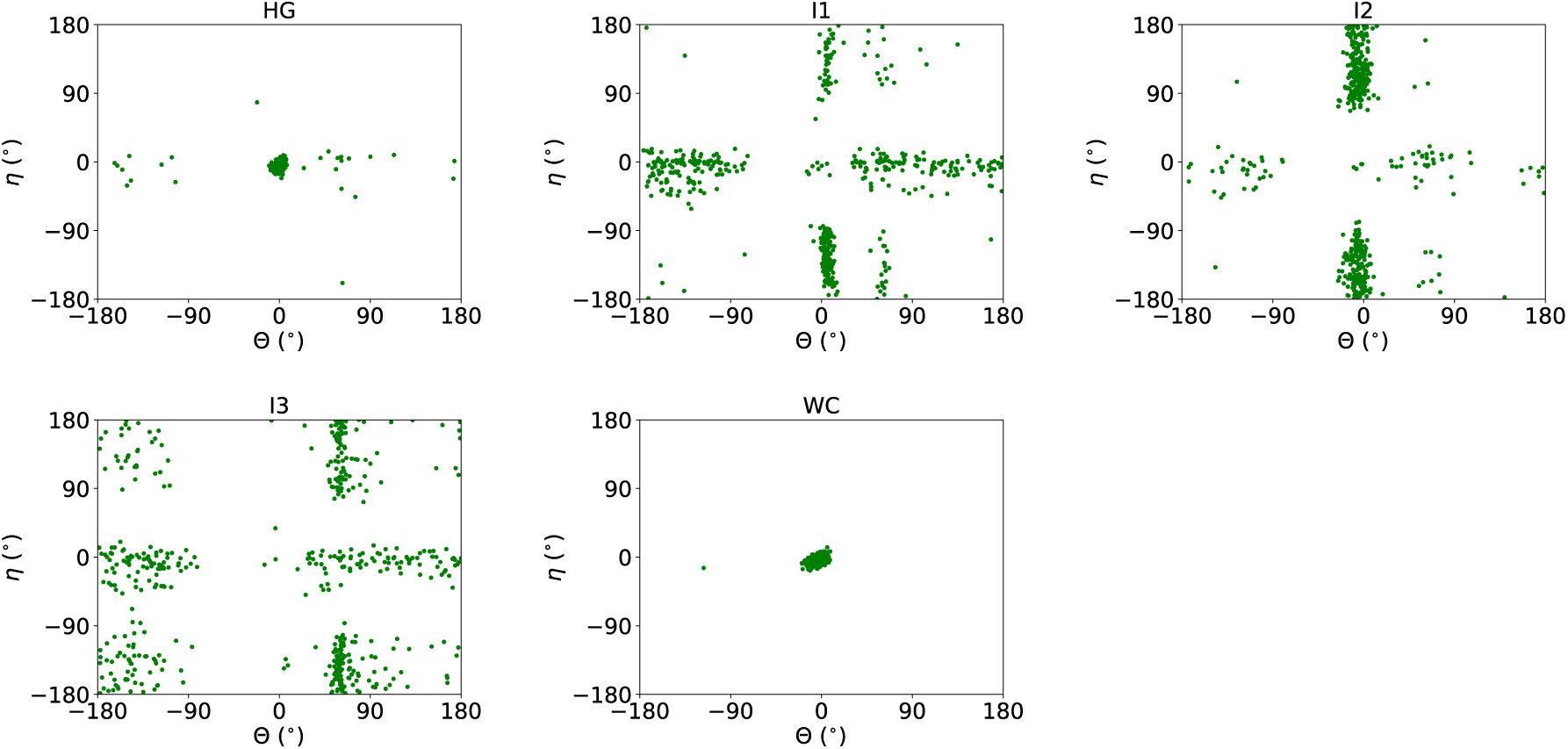
Distribution of pseudo dihedral angles of Adenine and Thymine base (Θ and **η** respectively) in the metastable states in A6-DNA

**Figure S11:**
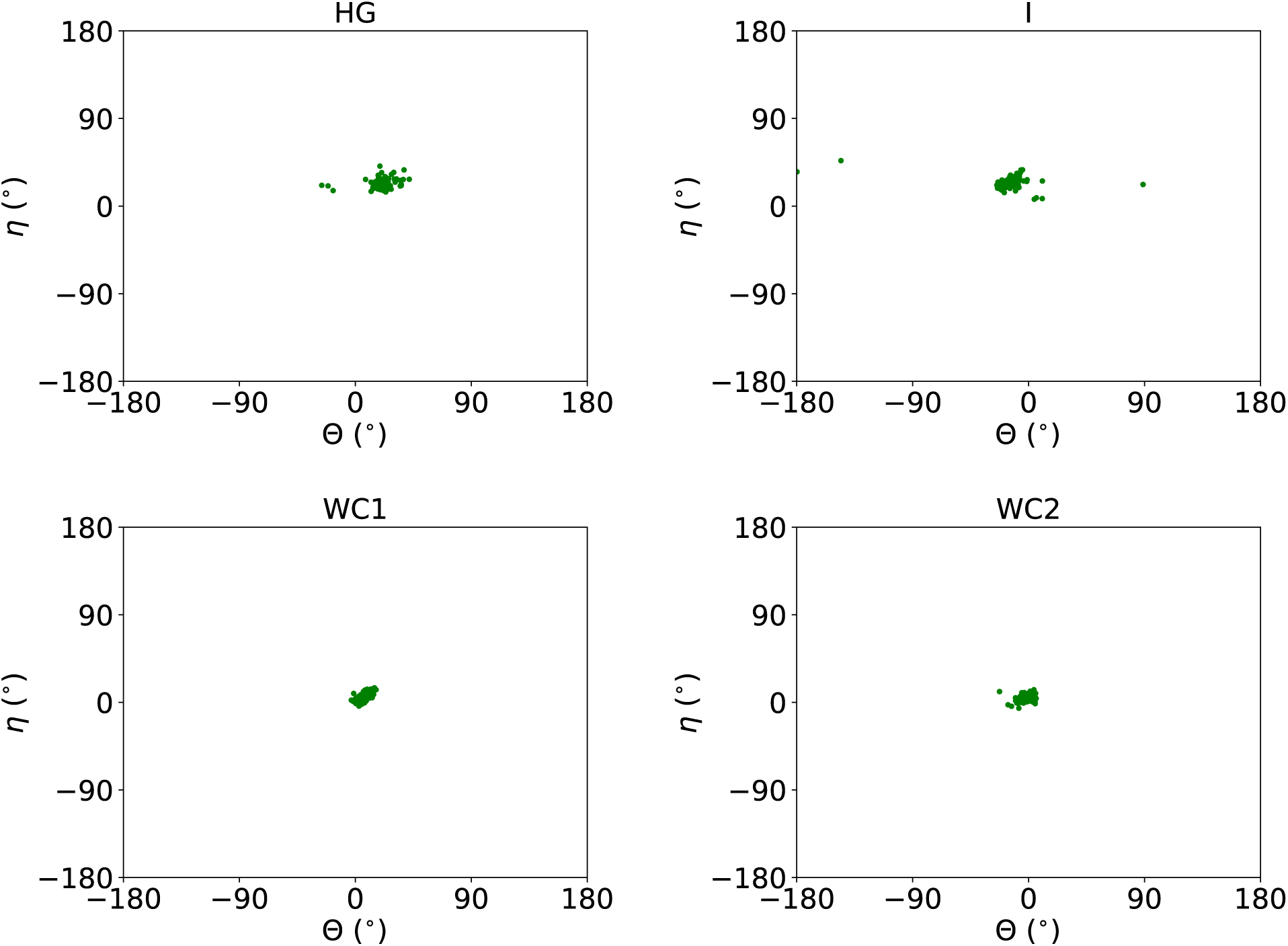
Same as Figure S10 but for A6-RNA.

